# BioMake: a GNU Make-compatible utility for declarative workflow management

**DOI:** 10.1101/093245

**Authors:** Ian H. Holmes, Christopher J. Mungall

## Abstract

The Unix “make” program is widely used in bioinformatics pipelines, but suffers from problems that limit its application to large analysis datasets. These include reliance on file modification times to determine whether a target is stale, lack of support for parallel execution on clusters, and restricted flexibility to extend the underlying logic program. We present BioMake, a make-like utility that is compatible with most features of GNU Make and adds support for popular cluster-based job-queue engines, MD5 signatures as an alternative to timestamps, and logic programming extensions in Prolog. BioMake is available from https://github.com/evoldoers/biomake under the Creative Commons Attribution 3.0 US license. The only dependency is SWI-Prolog, available from http://www.swi-prolog.org/. Contact: Ian Holmes or Chris Mungall.

## Introduction

The familiar Unix GNU Make utility has become a favored tool for “bioinformatics in-the-large” (Parker *et al.*, 2003). Alongside more elaborate workflow management systems, GNU Make holds its own for several reasons. Besides being ubiquitous and easy to use, with a simple syntax, it offers a powerful mix of *declarative logic* (the specification of target-dependency relationships from which Make deduces build chains) with *Unix scripting* (the lines of shell script that are executed when the build chain runs). GNU Make combines these elements with *functional programming*-inspired manipulation of text variables, lists, and directories, and includes Guile — GNU’s Scheme interpreter — as an extension language.

In our usage of GNU Make for data analysis, a common pattern is to analyze one or two examples manually, building up a `Makefile` recipe (or recipes), then scale the analysis up to the whole dataset. Makefiles remain, in our opinion, unrivalled for this purpose. However, GNU Make’s origins were as a tool for managing build pipelines, not large-scale data analyses, and it has several flaws that impede its use in bioinformatics.

## Results

We have developed a new tool, BioMake, that keeps the best features of GNU Make (including the ability to read a GNU `Makefile`) while addressing its shortcomings. Chief innovations of BioMake include:

### (1) MD5 signatures as an alternative to time-stamps

GNU Make uses file modification times to determine when files need to be rebuilt. This is fragile, especially on networked filesystems or cloud storage, where file timestamps may not be preserved or synchronized. In projects where a big data analysis can take hours or days, a spurious rebuild can be devastating, especially if it triggers further rebuilding of downstream targets. Instead of using timestamps, BioMake can be directed to use MD5 checksums: whenever a target is built, the MD5 hashes of that file and its dependents are recorded and stored. This can be used in combination with `Makefile` recipes that sort or canonicalize data to further guard against spurious rebuilds.

### (2) Support for cluster-based job queues

GNU Make can run multiple jobs in parallel, but only on one machine. It is possible to write cluster support directly into the `Makefile`, wrapping each recipe with a call to a job submission script, but this spoils GNU Make’s otherwise clean separation of concerns and often prevents it from tracking dependencies properly. BioMake has built-in support for Sun Grid Engine, PBS, and SLURM job submission systems, including dependency tracking (ensuring a target is not built until all its dependents have been built). It also (like GNU Make) offers built-in parallel execution on the same machine that BioMake is being run on.

### (3) Multiple wildcards per filename

GNU Make only allows a single wildcard (“stems”) in a filename, represented by the percent symbol (**%**) in the head of a recipe and by the automatic variable **$\*** in the body. In contrast, BioMake allows multiple wildcards: any unbound variable that appears in the head of a recipe can serve as a wildcard, and can subsequently be used in the body of the recipe.

### (4) Easy integration with ontologies and description logics

GNU Make’s domain-specific language extensions are based on Scheme, which is a functional language, but the underlying structure of a Makefile (rules such as “*to build A, you must first build* B” and “*to build B, you must first build C and D*”) is a logic program. BioMake’s domain-specific language is Prolog, making it trivially easy to incorporate ontologies and description logics such as the Gene Ontology (Blake, 2015) or the Sequence Feature Ontology (Eilbeck *et al.*, 2005). For example, we can easily create BioMake recipes for targets such as “*the whole-genome alignment for species X and Y, where X is a mammal and Y is a vertebrate*” or “*the GFF file containing co-ordinates of every human genomic feature of type T, where T is a term descended from ‘biological-region’ in the Sequence Ontology*”. In a Scheme program, we would have to write, test, and debug functions that explicitly generated these lists of terms and taxa; in a Prolog database, logical conditions such as “*X is a mammal*” or “*T is descended from biological-region*”are trivially easy to model directly, and the Prolog interpreter itself searches for all variable bindings that fit the model.

The Makefile in Figure 1 is a simple example illustrating multi-wildcard pattern-matching (point 3, above) and Prolog extensions (point 4). Such a Makefile could be used to build all alignment files whose names match the pattern `align-X-Y` where `X` and `Y` form an ordered pair of recognized species names.

**Figure 1:**
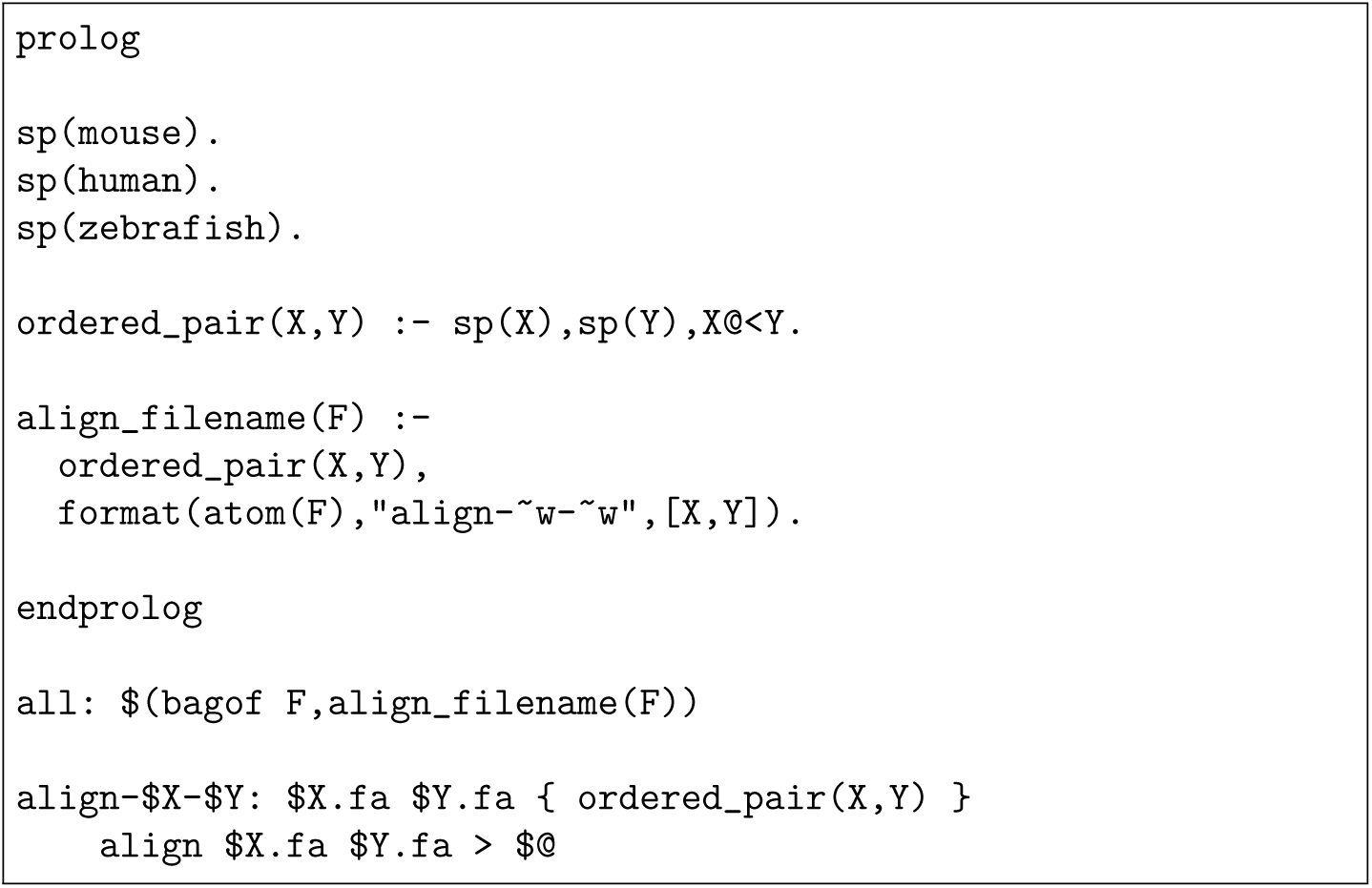
A hypothetical BioMake **Makefile** that runs **align** on all ordered pairs of files **mouse.fa**, **human.fa** and **zebrafish.fa**. The rule for file **align-$X-$Y** creates an alignment (using the program align, assumed to exist on the user’s **PATH**) from any two files **$X.fa** and **$Y.fa**. However, it only applied for those **$X** and **$Y** which are flagged as being valid species, via the Prolog facts **sp(X)** which appear between **prolog** and **endprolog** directives. The top-level target **all** uses BioMake’s **$(bagof…)** function, a wrapper for the Prolog predicate **bagof/3**, to find all ordered pairs of species that match the rule. This example is discussed in greater detail in the **README.md** file of the BioMake github repository.

## Discussion

We can contrast BioMake with other solutions for bioinformatics workflow management. Systems such as CWL^1^ or Galaxy^2^ have many useful features such as web interfaces and cloud support, but they do not deduce the workflow from a series of declaratively-specified dependencies: rather, they require explicit specification of inputs, outputs, and ordering of tasks.

There are also other GNU Make clones and variants, including some in functional/parallel languages (Erlang make^3^), offering MD5 signatures (omake^4^, makepp^5^) or supporting cluster-based parallelism (e.g. Oracle Grid Engine’s qmake^6^). Some of these overlap in functionality with BioMake, but none offers the full feature set described here.

Prolog for bioinformatics may seem esoteric, but it is a good fit for some applications; both of us have used it before. Blipkit is a Prolog toolkit for logic programming on ontologies and other data structures (Mungall, 2009). PRISM is a probabilistic dialect of Prolog that was used to implement HMMs, trellis models, and other probabilistic modeling for sequence annotation (Mørk and Holmes, 2012; Have and Mørk, 2014). BioMake is complementary to these applications, as it can stand alone as a workflow management tool, but can also be integrated with other Prolog approaches for additional power.

## Footnotes

1 http://commonwl.org/, accessed Dec 9, 2016.

2 https://usegalaxy.org/, accessed Dec 9, 2016.

3 http://erlang.org/doc/man/make.html, accessed Dec 9, 2016.

4 http://omake.metaprl.org/, accessed Dec 9, 2016.

5 http://makepp.sourceforge.net/, accessed Dec 9, 2016.

6 http://gridscheduler.sourceforge.net/htmlman/htmlman1/qmake.html, accessed Dec 9, 2016.

